# Population mobility and dengue fever transmission in a major city in Southeastern Brazil, 2007-2015

**DOI:** 10.1101/825109

**Authors:** Igor C. Johansen, Marcia C. Castro, Luciana C. Alves, Roberto L. Carmo

## Abstract

**Background:** Around 14% of world dengue virus (DENV) cases occur in the Americas, the majority of them in Brazil. Although socioeconomic, environmental and behavioral correlates of dengue have been analyzed for different contexts, the role played by population mobility on DENV epidemics, especially at the local level, remains scant. This study assesses whether the daily pattern of population mobility is associated with DENV transmission in Campinas, a Brazilian major city with over 1.2 million inhabitants in São Paulo state.

**Methodology/Principal Findings:** DENV notifications from 2007 to 2015 were geocoded at street level (n=114,884) and combined with sociodemographic and environmental data from the 2010 Population Census. Population mobility was extracted from the Origin-Destination Survey (ODS), carried out in 2011, and daily precipitation was obtained from satellite imagery. Zero-Inflated Negative Binomial (ZINB) regression models controlled by demographic and environmental factors revealed that high population mobility had a substantial positive effect on higher risk for DENV transmission. High income and residence in apartments were found to be protective against the disease, while unpaved streets, number of strategic points (such as scrapyards and tire repair shops), and precipitation were consistently risk factors for DENV infection.

**Conclusions/Significance:** The use of fine-scale geographical data can unravel transmission idiosyncrasies not evident from a coarse spatial analysis. Even in a major city like Campinas, the vast majority of population daily mobility occurs at short distances. Based on our results, public policies on DENV transmission control should dedicate special attention to local hubs of population mobility, especially during high transmission weeks and in high dengue incidence areas.

**Author Summary:** Currently, about half of the world population is at risk of a dengue infection. Numerous studies have addressed the socioeconomic and environmental determinants of the disease. However, little is known about the role played by population mobility on dengue transmission, particularly at the local scale. This study aims at investigating this issue. Our hypothesis was that population movements are a prominent driving force for dengue diffusion locally. We investigated the case of Campinas, a municipality with over 1.2 million inhabitants in Brazil that recorded dengue epidemics in 2007, 2014 and 2015. Our study focused on the years 2007 to 2015, comprising more than 114 thousand cases, geocoded to the household address, and combined with socioeconomic, environmental and daily population mobility data. Our results showed that even controlling for demographic and environmental factors, population mobility was the most important predictor for dengue fever incidence.

## Introduction

Despite the growing concern about other infectious diseases transmitted by *Aedes aegypti*, such as Zika virus and chikungunya, dengue virus (DENV) remains a global threat [1–3]. The incidence of this disease has grown dramatically in recent decades. Economically, the annual global cost of DENV is estimated at around US$8.9 billion [4] and nearly 390 million dengue infections occur per year, of which 96 million manifest clinically [5]. About 3.9 billion people in 128 countries are at risk of infection [6]. Regionally, approximately 70% of dengue cases are observed in Asia, followed by Africa (16%) and the Americas, 14% [5]. In 2016, more than 2.3 million DENV cases were reported in the Americas, 64% of them in Brazil [7].

Historically, Brazil succeeded in eliminating *Ae. aegypti* during the 1940s, and that campaign inspired the Pan American Health Organization (PAHO) to pursue elimination in the Americas in the 1950s [8]. Although *Ae. aegypti* was eliminated from the Americas (with the exception of Colombia, Venezuela, British Guiana, Surinam, and the United States), the mosquito was reestablished in the early 1970s [3]. The reintroduction of DENV in Brazil occurred at the beginning of the 1980s, through the northern state Amapá [9]. Since then, the disease has grown from an incidence rate close to zero in 1985 to more than 830 cases per 100,000 inhabitants in 2015 [10]. In 2019 (until April), DENV incidence rate reached 131 cases per 100,000 inhabitants [11].

In Brazil’s most populous state, São Paulo, DENV transmission began to be reported, with clinical and laboratory diagnosis, in March 1984 [12]. Campinas is the third most populous city in São Paulo, having first reported DENV autochthonous transmission in 1996 [13]. Since then the disease remained endemic, with major epidemics recorded in 2007, 2014 and 2015. Preliminary 2019 data suggest that a new epidemic is in course in the municipality [11]. Therefore, DENV remains a public health challenge in Campinas.

While socioeconomic, environmental, and behavioral correlates of dengue have been analyzed for different contexts in Brazil [14–17], more information is needed to comprehend the role played by population mobility, especially at the local level. This study addresses this issue, and our hypothesis is that population movements are a prominent driving force for dengue diffusion locally. The analysis considered nine consecutive years (2007 to 2015), addressing temporal and spatial variations.

We chose Campinas because (i) dengue is an increasing public health threat in the city; (ii) in 2014, Campinas registered the highest number of DENV cases in the Brazil (7% of the total cases, while it shares only 0.6% of population in the country); (iii) in 2015, it recorded the most prominent DENV incidence rate among the municipalities with over 1 million inhabitants; (iv) it is highly diverse in socioeconomic status of population groups and in access to urban infrastructure (social inequalities), being similar to other Brazilian and Latin American urban contexts; and, most importantly, (v) Campinas has collected unique data on regular daily mobility.

## Materials and Methods

### Study area

The municipality of Campinas (22°53’20”S, 47°04’40”W) is located in Southeastern Brazil, the most economically dynamic region of the country. It has approximately 1.2 million inhabitants and is 100 km from the city of São Paulo, the capital of São Paulo state [18]. Campinas is the geographical center and main city of the Campinas Metropolitan Region, composed of 20 municipalities, with 2.8 million inhabitants [19]. With an area of around 800 km^2^, the space occupation in Campinas municipality is highly heterogeneous. Although population density is about 1,360 people per km^2^, it considerably varies from highly populated to nearly empty neighborhoods. The weather is tropical, with rains during the summer (December to March), and drought in the winter (June to September). The average minimum temperature is 19°C (66.2F), and the maximum is 29°C (84.2F).

Despite its location within an economically developed area of the country, the city has patent contrasts in terms of population socioeconomic characteristics and urban infrastructure. There are still gaps in access to water, sewage, and garbage collection. Although the 2010 Population Census [20] reports that almost all the population had access to piped water (99%), the regularity of the supply can vary according to the area of the city, a problem intensified during the severe drought the city faced in 2014, with subsequent water shortages [21]. Garbage and sewage collection are also virtually widespread, reaching 99% and 87% of the population, respectively [20], although the remaining gaps are greater especially in the outskirts of the city.

The most affluent groups live in the center and north of the municipality, where urban infrastructure is of better quality. In contrast, the south concentrates the impoverished people, with less access to urban services. The urban configuration of Campinas encourages population mobility, given that the territory is abundantly crossed by major roads and highways (Fig 1).

**Fig 1.**
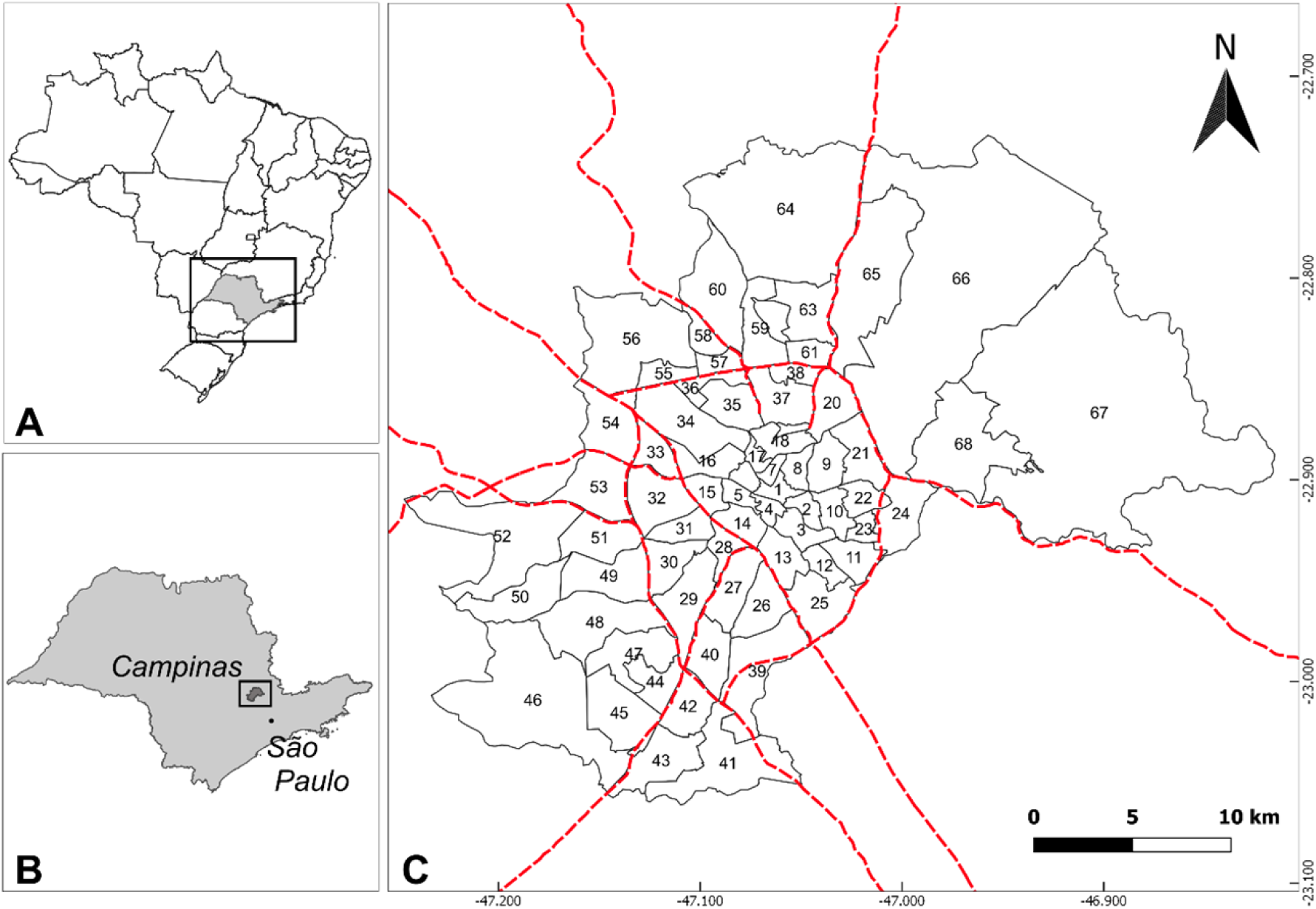
Origin-Destination (OD) units of Campinas, São Paulo, Brazil. A: Brazil state boundaries; B: São Paulo state; C: Campinas municipality. Red dotted lines refer to the main state roads that cross Campinas. Numbers refer to OD units. Unit 1 (center of the figure) comprises the city center. An OD name key is included in the supplementary material (S1 Table). Figure created with QGIS software version 3.8, an open source Geographic Information System (GIS) licensed under the GNU General Public License (https://qgis.org/en/site/about/index.html). Publicly available shapefiles provided from the Brazilian Institute of Geography and Statistics (IBGE) website (https://www.ibge.gov.br/geociencias/downloads-geociencias.html).

### Dengue transmission in Campinas

Campinas had its first large dengue epidemic in 2007, with 1,100 cases per 100,000 inhabitants. Two consecutive epidemics followed in 2014, with 3,647 cases per 100,000 inhabitants, and in 2015, when over 5,600 cases per 100,000 inhabitants were registered. Each epidemic cycle corresponded to the circulation of a new serotype: DEN-3 in 2007, and DEN-1 in 2014 and 2015 [22].

### Variables and data sources

We studied 114,884 reported DENV autochthonous cases from 1^st^ January 2007 to 31^st^ December 2015, provided by the Notifiable Diseases Information System [23]. The spatial location of the cases was recorded at patients’ residence addresses, and the temporal detail was based on the date of the beginning of symptoms. Following protocols from the Brazilian Ministry of Health [24], the majority of cases were diagnosed using the clinical-epidemiological criteria, and a small number of cases were analyzed using serologic exams and virus isolation, which occurs especially for more severe cases and deaths.

To assess if high mobility was associated with high dengue incidence, while controlling for other correlates, we used information about population mobility in Campinas obtained from the Origin-Destination Survey (ODS), carried out in 2011 [25]. ODS included all population travels between or within Origin Destination Survey units (from now on called only OD units) during one regular business day (Monday to Friday); interviews were carried out from September to November 2011. The assumption in using these data was that DENV infected patients are mobile.

ODS divided Campinas into 68 units, based on the transportation zoning system, urban equipment infrastructure, and physical barriers. In this study, we used 66 OD units as the unit of analysis (two units were not inhabited). The reasons for using the OD units were two-fold: (i) they comprised two or more Census tracts, making it straightforward to merge mobility with Census data; and (ii) households were sampled within each area, impeding the reorganization of this information on other spatial units under the risk of biasing the survey results.

For each of the 66 OD units, we calculated dengue incidence rates (cases per 100,000 inhabitants) over the 469 epidemiological weeks of the study period. The epidemiological week is the main temporal unit that the Brazilian Ministry of Health uses to organize, process and plan health policies in the country, including those concerning infectious diseases.

Additionally, we calculated covariates that could influence DENV transmission. These covariates can broadly be categorized as demographic variables reflecting peoples’ attributes (household per capita income, sex ratio, and population density), and environmental variables (residence in apartments, unpaved streets, and open sewage). These variables were extracted from the 2010 Population Census [20] and aggregated to the OD unit level.

We also gathered data on Strategic Points (SPs), comprising locations such as junkyards, tire repair shops, deposits of recyclable materials, etc. They are essential for understanding dengue occurrence as these areas present huge potential to accumulate *Aedes* breeding sites [24,26,27]. Examples are shown in Fig 2. The addresses of SPs in Campinas were obtained from the Superintendence for Control of Endemic Diseases [28], which allowed georeferencing and then obtaining the total number of SPs per OD unit. Although the information on SP locations was available only for the year 2017, our assumption was that SPs can change from one specific location to another, but their relative spatial distribution within Campinas tends to be similar over time.

**Fig 2.**
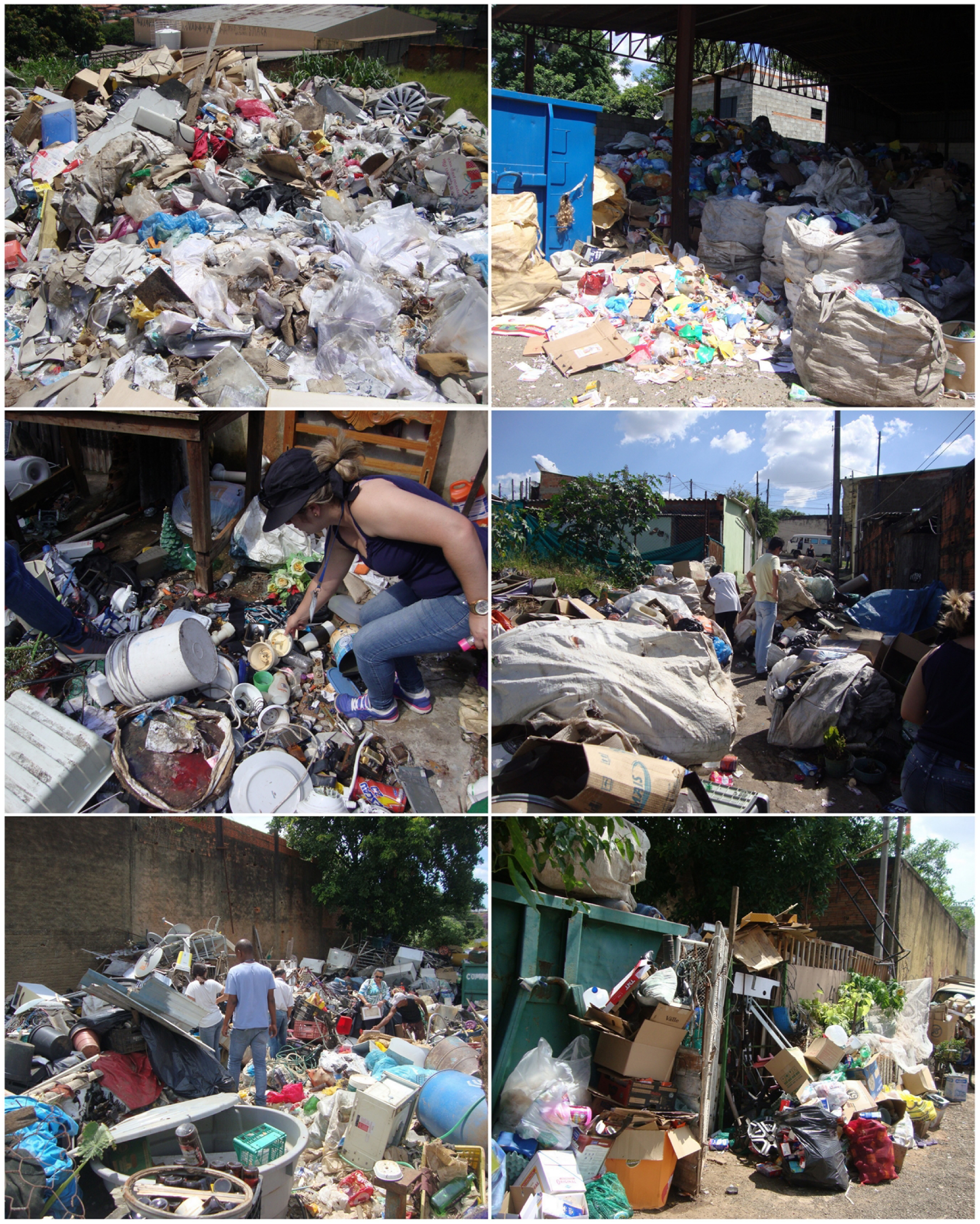
Strategic Points in Campinas. Pictures were taken during fieldwork (Jan. 2018).

Concerning environmental factors, we compiled data about rainfall, given water is essential for the vector to breed. Weekly information on precipitation was extracted from the Climate Hazards Group InfraRed Precipitation with Station data [29]. This dataset provides the estimated precipitation per day based on satellite imagery, at a 5 Km resolution. The accumulated precipitation was calculated per epidemiological week for each OD unit. After an assessment of previous works [17,26,30–32], we tested different lags: 2 weeks, 4 weeks (1 month), 8 weeks (2 months), and 12 weeks (3 months). This lag aimed to cover the expected time necessary for the precipitating water to pool, and for the entire cycle of transmission (from oviposition to a confirmed dengue case) to take place [17,33,34].

To better capture the intensity of transmission and of mobility, three categorical variables were considered. The first, “high week”, comprised weeks containing a proportion of dengue cases higher than 1.52% of total annual cases, which would be expected if cases were equally distributed along the weeks of the year (0-No, 1-Yes), as shown in S1 Fig. The second, “high area”, distinguished the ODs with more than 300 cases per 100,000 inhabitants, per week (0-No, 1-Yes). This threshold was chosen inspired by the Brazilian standard criteria to assess dengue incidence rates that are calculated annually [24]; our study used the same criteria, but considering the week, in order to highlight the most prominent areas with high DENV figures. Finally, the variable “high mobility” identified areas with population mobility higher than the median observed in the city: 1,800 travels per 1,000 inhabitants (0-No, 1-Yes); it was calculated considering the differences in mobility levels between different ODs.

### Analytical approach

A Zero-Inflated Negative Binomial regression (ZINB) model was used. The ZINB model accounts for extra-variation (overdispersion) in the data [35–37]. The outcome variable was dengue incidence rate: DENV cases per 100,000 inhabitants per OD unit and epidemiological week. Covariates included demographic (household per capita income, sex ratio, population density) and environmental factors (residence in apartments, unpaved streets, open sewage, number of strategic points, and accumulated precipitation). Additionally, dummy variables were used for high mobility, high epidemiological weeks and high incidence areas. The selection of independent variables to include in the final models was based on the stepwise backward process. The final model was chosen based on Akaike Information Criterion (AIC) for alternative distributions, such as negative binomial, Poisson, and zero-inflated.

Considering the variation of DENV distribution through space and time, we run five models. Model 1 included all observations (N=30,954 OD-weeks). Models 2 and 3 were stratified by high (N=8,778 OD-weeks) and low epidemiological weeks (N=22,176 OD-weeks). Models 4 and 5 stratified the analysis by intensity of transmission, the former considering only high areas (N=637 OD-weeks) and the latter the low areas (N=30,317 OD-weeks). Models were controlled by year, taking 2015 as reference, accounting for differences between yearly variation in incidence rates. In all of these models, the variable population mobility was exactly the same, and statistical significance was evaluated at a p-value < 0.01.

Geographical Information System (GIS) tools for merging different datasets were carried out in ArcGIS 10.5.1. Data cleaning, descriptive and statistical analyses were performed in Stata 14.0 (Stata Corp., College Station, TX, USA).

### Ethics statement

This research does not require an Institutional Review Board approval because it comprises an analysis of secondary and anonymized data.

## Results

Between 2007 and 2015, 123,042 dengue cases were reported in Campinas, ranging from 159 in 2009 to 58,720 in 2015. In total, 114,884 (93.4%) had sufficient information to be geocoded at the household level and included in the analysis. Geocoding success was high, and varied from 90.0% in 2014 to 99.4% in 2009 (Table 1).

**Table 1.** Number of reported dengue cases and total successfully geocoded by year, Campinas, 2007-2015.

| Year | Reported dengue cases (A) | Geocoded (B) | % (B/A) |
| --- | --- | --- | --- |
| 2007 | 10,385 | 10,195 | 98.2 |
| 2008 | 267 | 265 | 99.3 |
| 2009 | 159 | 158 | 99.4 |
| 2010 | 2,131 | 2,097 | 98.4 |
| 2011 | 2,783 | 2,746 | 98.7 |
| 2012 | 534 | 513 | 96.1 |
| 2013 | 6,344 | 5,769 | 90.9 |
| 2014 | 41,719 | 37,564 | 90.0 |
| 2015 | 58,720 | 55,577 | 94.6 |
| Total | 123,042 | 114,884 | 93.4 |

Table 2 presents descriptive statistics of the continuous OD-level variables. DENV incidence rate per 100,000 inhabitants by OD-weeks, ranged from the mean 0.21 in 2009 to 96.92 in 2015. The smallest standard deviation was observed in 2009, 1.93, while the highest was 256.30, in 2015. After investigating different time lags: 2, 4, 8 and 12 weeks, we found that the 8-weeks lag (two months) presented the highest coefficient associated with the dependent variable (dengue incidence rate). The sum of dengue cases and the accumulated precipitation lagged 8 weeks is presented in S2 Fig. The accumulated precipitation (lagged by 8 weeks) ranged from an average of 24.9 mm in 2014 to 32.3 mm in 2012. Mean household income per capita by OD was R$ 1,347.13, and the total population mean sex ratio among OD units was 95.46. Population density was, on average, 3,260 persons per km^2^, although it considerably varied from the minimum 15 to the maximum 13,180. OD areas presented an average of 9.8 SPs, ranging from 0 to 65. Also, around 19% of all households were classified as apartment, these types of residence were most common in the city center (up to 95%). On average, 7.21% of households in the municipality were located in unpaved streets, varying substantially from one location to the other: from 0% in well-provided OD units to nearly 60% of unpaved streets in others. Also, an average of 3.82% households presented open sewage, ranging from 0% to more than 38% across the different units of analysis.

**Table 2.** Descriptive statistics of the continuous OD-level variables.

| Variable |  | Mean | SD | Range |
| --- | --- | --- | --- | --- |
| **Reported dengue cases by OD-week (outcome)** |  |  |  |  |
| **Count (number of cases)** | 2007 | 2.97 | 8.88 | (0-97) |
|  | 2008 | 0.77 | 0.33 | (0-6) |
|  | 2009 | 0.46 | 0.28 | (0-5) |
|  | 2010 | 0.61 | 2.16 | (0-43) |
|  | 2011 | 0.80 | 2.65 | (0-35) |
|  | 2012 | 0.15 | 0.60 | (0-11) |
|  | 2013 | 1.68 | 6.20 | (0-106) |
|  | 2014 | 10.74 | 33.91 | (0-481) |
|  | 2015 | 16.19 | 47.09 | (0-527) |
| **Rate (cases per 100,000)** | 2007 | 17.99 | 66.88 | (0-1,498.1) |
|  | 2008 | 0.60 | 7.80 | (0-3,74.5) |
|  | 2009 | 0.21 | 1.93 | (0-87.1) |
|  | 2010 | 3.79 | 16.58 | (0-374.5) |
|  | 2011 | 6.03 | 34.92 | (0-1,084.0) |
|  | 2012 | 0.91 | 8.20 | (0-374.5) |
|  | 2013 | 10.22 | 40.66 | (0-749.1) |
|  | 2014 | 63.22 | 164.66 | (0-2,264.8) |
|  | 2015 | 96.92 | 256.30 | (0-4,494.4) |
| **Climate (exposure)** |  |  |  |  |
| Precipitation<br>Lagged 8 weeks<br>(sum in mm) | 2007 | 27.31 | 33.53 | (0-188.1) |
|  | 2008 | 28.86 | 26.32 | (0-137.56) |
|  | 2009 | 26.09 | 26.31 | (0-109.4) |
|  | 2010 | 30.19 | 37.78 | (0-163.8) |
|  | 2011 | 28.91 | 36.62 | (0-208.9) |
|  | 2012 | 32.32 | 33.68 | (0-172.5) |
|  | 2013 | 32.09 | 31.58 | (0-120.5) |
|  | 2014 | 24.90 | 25.87 | (0-124.1) |
|  | 2015 | 28.09 | 26.25 | (0-108.7) |
| **Demographic (exposure)** |  |  |  |  |
| Household per capita income (R\$) | | 1,347.13 | 850.18 | (397.4-4,031.8) |
| Total pop. sex ratio (male/female per 100) |  | 95.46 | 8.15 | (78.7-118.7) |
| Population density (persons/km<sup>2</sup>) |  | 3,260.38 | 2,878.71 | (15.1-13,180.3) |
| **Environmental (exposure)** |  |  |  |  |
| Number of Strategic Points (SPs) |  | 9.8 | 13.6 | (0-65) |
| Residence in apartments (%) |  | 18.94 | 22.46 | (0-95.0) |
| Households in unpaved streets (%) |  | 7.21 | 11.34 | (0-59.8) |
| Households with open sewage (%) |  | 3.82 | 8.1 | (0-38.4) |

With regards to the intensity of transmission and of mobility (Table 3), there were 8,778 OD-weeks (28.4% of the total) with higher proportion of dengue cases than expected, while 637 OD-weeks were classified as High Areas (2.1% of the total), i.e. with elevated incidence rates. Additionally, given the variable High Mobility uses the median to divide the OD areas in terms of population mobility, areas with high and low mobility are numerically identical, with 15,477 OD-weeks in each (50%).

**Table 3.** Dummy variable definitions and descriptive statistics.

| Variable and Description | Value | Number of observations (%) | Total |
| --- | --- | --- | --- |
| **Scale (exposure)** |  |  |  |
| High Week (proportion of cases > 1.52% of the annual cases) | 0 = No<br>1 = Yes | 22,176 (71.6)<br>8,778 (28.4) | 30,954 |
| High Area (> 300 cases per 100,000 inhabitants, by epidemiological week) | 0 = No<br>1 = Yes | 30,317 (97.9)<br>637 (2.1) | 30,954 |
| **Mobility (exposure)** |  |  |  |
| High Mobility (population mobility > 1,800 median travels per 1,000 inhabitants) | 0 = No<br>1 = Yes | 15,477 (50)<br>15,477 (50) | 30,954 |

The spatial distribution of the total number of epidemiological weeks with high incidence rates (Fig 3) shows a concentration of “High Areas” in the North of the municipality, encompassing neighborhoods such as Cidade Universitária, Real Parque, Jardim Santa Izabel, Parque das Universidades, Jardim São Marcos, and Jardim Santa Mônica. It also highlights the importance of DENV cases in the Southwest region, where we find Ouro Verde, Satélite Íris, Florence, Cosmos, Sirius, Jardim Paulicéia, and Jardim Campos Elísios neighborhoods. These areas are the ones with the highest historical concentration of dengue cases in the city during the study period, and where epidemics recurrently occur (Fig 1 and S1 Table show the location of each OD unit and the neighborhoods comprised under each OD unit, respectively).

**Fig 3.**
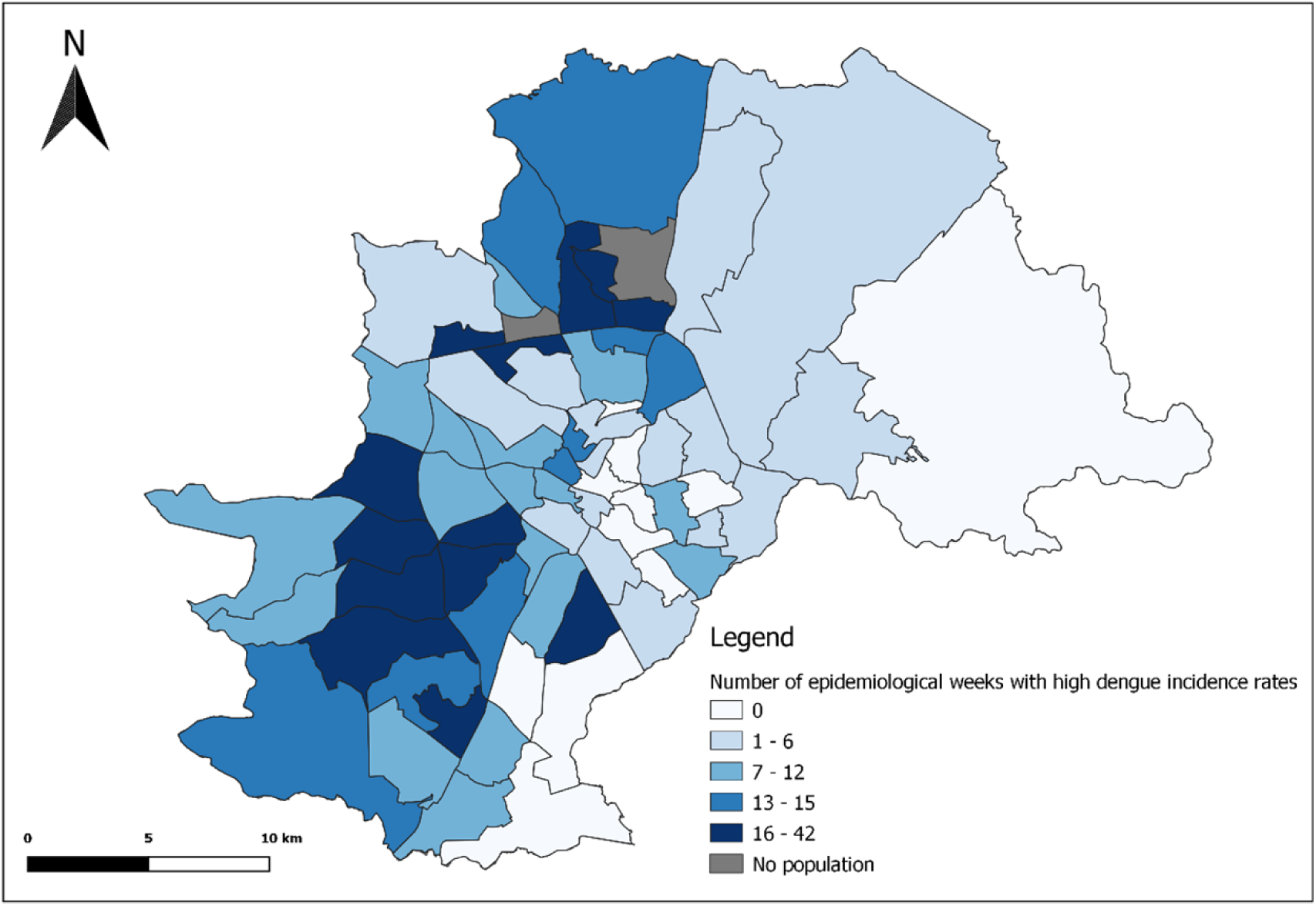
“High Areas” - Sum of epidemiological weeks with high dengue incidence rates (>300 cases per 100,000 persons), Campinas, 2007-2015. Variation represented in shades of blue, from areas with no epidemiological week with high dengue incidence rates to areas that registered the peak of 16 to 42 epidemiological weeks with high incidence rates during the study period. Figure created with QGIS software version 3.8, an open source Geographic Information System (GIS) licensed under the GNU General Public License (https://qgis.org/en/site/about/index.html). Publicly available shapefiles provided from the Brazilian Institute of Geography and Statistics (IBGE) website (https://www.ibge.gov.br/geociencias/downloads-geociencias.html). Data on dengue fever cases are publicly available under the Brazilian “Information Access Law” (12.527/2011), obtained from the Notifiable Diseases Information System (SINAN).

As for the intensity of mobility, while the “most mobile” populations were apparently widespread in Campinas, they mostly overlapped with “High Areas” distribution, which beforehand suggested a link between population mobility and dengue occurrence, as shown in Fig 4. The exception to this pattern is the city center, in which we notice high mobility but no expressive records of dengue occurrence.

**Fig 4.**
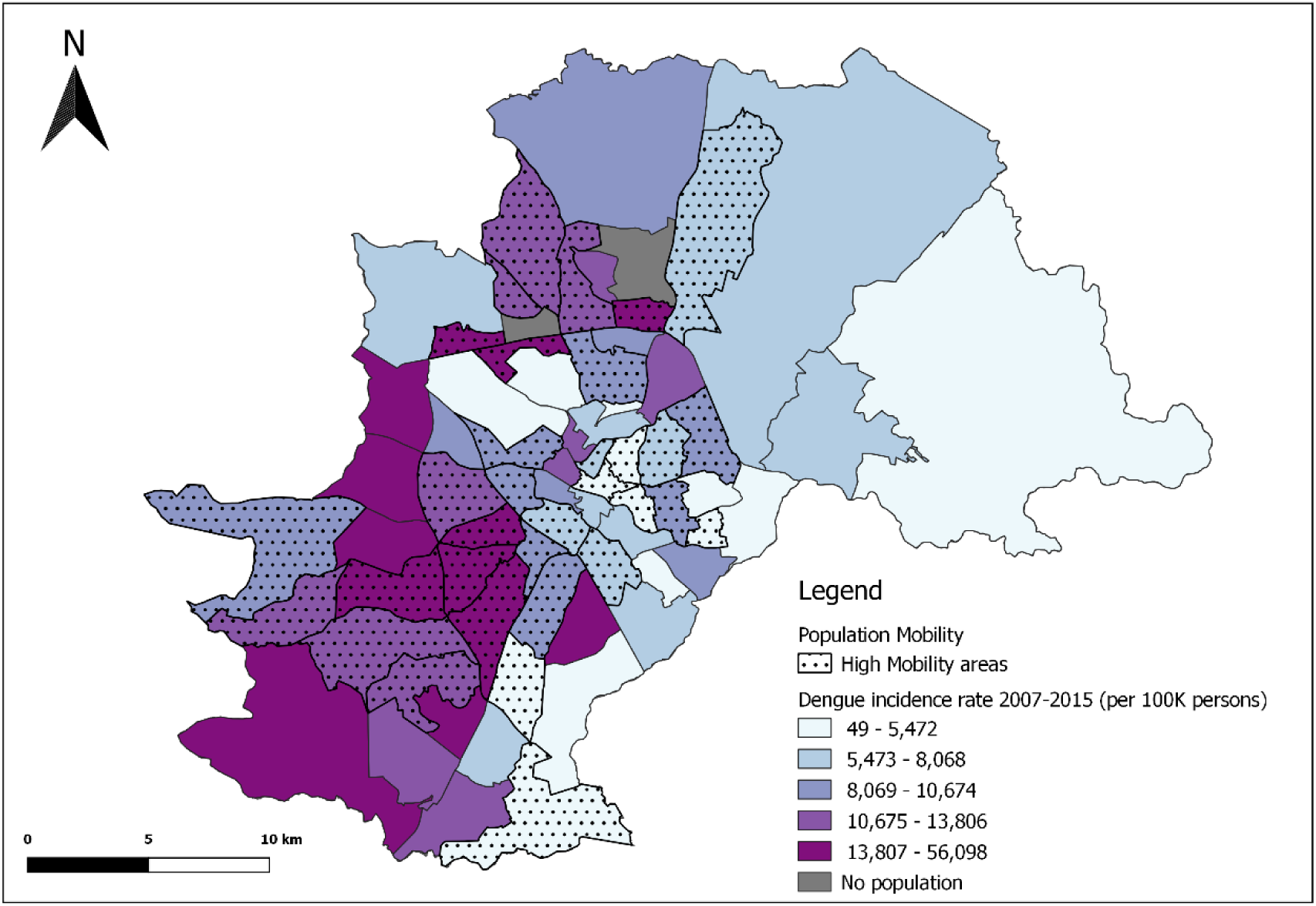
“High Mobility” - Areas with population mobility higher than the 1,800 travels per 1,000 inhabitants and dengue incidence rate (per 100,000 persons), Campinas, 2011 and 2007-2015 respectively. Dotted areas represent units of analysis with high mobility. Dengue incidence rates classified in 5 categories ranging from 49 to more than 56,000 cases per 100,000 persons, colored from light blue (lowest) to dark purple (highest incidence rates). Figure created with QGIS software version 3.8, an open source Geographic Information System (GIS) licensed under the GNU General Public License (https://qgis.org/en/site/about/index.html). Publicly available shapefiles provided from the Brazilian Institute of Geography and Statistics (IBGE) website (https://www.ibge.gov.br/geociencias/downloads-geociencias.html). Data on dengue fever cases are publicly available under the Brazilian “Information Access Law” (12.527/2011), obtained from the Notifiable Diseases Information System (SINAN).

In assessing the role of population mobility, the complete model (Model 1, Table 4), showed that, on average, living in a highly mobile area was associated with an increase of 40% in dengue incidence rates. Over the entire study period, the increase of one SP in a study unit was accompanied by an increment, on average, of 3% in dengue incidence rates. Similarly, the increase of 1% in unpaved roads and open sewage was associated with an increase of 1% in dengue incidence rates. Residence in apartments, on the other hand, was protective against the disease, although barely significant. Indicator variables High Week and High Area were statistically significant and presented coefficients with high magnitude, supporting stratified analyses.

**Table 4.** ZINB model considering all OD-weeks (Model 1).

| Parameter |  | Model 1 (n=30,954) |  |  |  |
| --- | --- | --- | --- | --- | --- |
|  |  | IRR | 95% CI | z | p-value |
| Climate | Precipitation | 1.01 | [1.00-1.01] | 13.74 | <0.01 |
| Demographic | Income | 0.99 | [0.99-0.99] | -6.11 | <0.01 |
|  | Sex Ratio | 0.94 | [0.94-0.95] | -13.71 | <0.01 |
|  | Pop. Density | 1.00 | [1.00-1.00] | 13.84 | <0.01 |
|  | High Mob. | 1.41 | [1.32-1.49] | 10.96 | <0.01 |
| Environmental | Strat. Points | 1.03 | [1.02-1.03] | 22.13 | <0.01 |
|  | Apartment | 0.98 | [0.98-0.99] | -13.82 | <0.01 |
|  | Unpaved | 1.01 | [1.01-1.01] | 5.56 | <0.01 |
|  | Open Sewage | 1.01 | [1.01-1.02] | 4.51 | <0.01 |
| Scale | High Week | 15.19 | [14.32-16.11] | 90.24 | <0.01 |
|  | High Area | 3.01 | [2.67-3.41] | 17.57 | <0.01 |
| Year<br>*Ref.: 2015* | 2007 | 0.21 | [0.19-0.23] | -35.14 | <0.01 |
|  | 2008 | 0.01 | [0.01-0.01] | -57.63 | <0.01 |
|  | 2009 | 0.00 | [0.00-0.00] | -55.66 | <0.01 |
|  | 2010 | 0.05 | [0.04-0.05] | -55.87 | <0.01 |
|  | 2011 | 0.05 | [0.05-0.06] | -56.22 | <0.01 |
|  | 2012 | 0.01 | [0.01-0.01] | -64.93 | <0.01 |
|  | 2013 | 0.11 | [0.10-0.12] | -45.59 | <0.01 |
|  | 2014 | 0.64 | [0.59-0.69] | -11.01 | <0.01 |

Tables 5 and 6 show stratified models by high and low epidemiological weeks (Models 2 and 3), and Tables 7 and 8 present stratified models by high and low incidence areas (Models 4 and 5). Results indicate that precipitation was positively associated with dengue incidence rates in periods of low transmission weeks and low incidence areas (Models 3 and 5). The variable households in unpaved streets was significantly correlated with dengue incidence rates in high transmission weeks and areas (Models 2 and 4). Living in high mobility areas was consistently a risk a factor for dengue incidence rates across all models. On average, living in a highly mobile area increased dengue incidence rate in 46% and 69% during high transmission weeks and in high transmission areas, respectively (Models 2 and 4). During low transmission weeks and in low transmission areas, being part of a highly mobile population increased by 26% and 35% dengue incidence rates, respectively (Models 3 and 5). Similarly to the complete model (Model 1), the mobility variable had the largest association with dengue incidence across all stratified models.

**Table 5.** ZINB model considering data stratified by High Weeks (Model 2).

| Parameter |  | Model 2 (n=8,778) |  |  |  |
| --- | --- | --- | --- | --- | --- |
|  |  | IRR | 95% CI | z | p-value |
| Climate | Precipitation | 0.99 | [0.99-1.00] | -1.61 | 0.107 |
| Demographic | Income | 0.99 | [0.99-0.99] | -9.14 | <0.01 |
|  | Sex Ratio | 0.93 | [0.92-0.93] | -19.48 | <0.01 |
|  | Pop. Density | 1.00 | [1.00-1.00] | 15.86 | <0.01 |
|  | High Mob. | 1.46 | [1.38-1.55] | 12.71 | <0.01 |
| Environmental | Strat. Points | 1.03 | [1.02-1.03] | 21.20 | <0.01 |
|  | Apartment | 0.98 | [0.98-0.98] | -15.87 | <0.01 |
|  | Unpaved | 1.02 | [1.01-1.02] | 8.70 | <0.01 |
|  | Open Sewage | 1.01 | [1.00-1.01] | 2.85 | <0.01 |
| Scale | High Area | 3.10 | [2.85-3.38] | 26.24 | <0.01 |
| Year<br>*Ref.: 2015* | 2007 | 0.29 | [0.27-0.33] | -25.76 | <0.01 |
|  | 2008 | 0.01 | [0.00-0.01] | -56.26 | <0.01 |
|  | 2009 | 0.00 | [0.00-0.00] | -53.78 | <0.01 |
|  | 2010 | 0.07 | [0.06-0.07] | -51.16 | <0.01 |
|  | 2011 | 0.07 | [0.06-0.08] | -52.58 | <0.01 |
|  | 2012 | 0.01 | [0.01-0.01] | -67.91 | <0.01 |
|  | 2013 | 0.15 | [0.14-0.17] | -38.41 | <0.01 |
|  | 2014 | 0.84 | [0.77-0.91] | -4.13 | <0.01 |

**Table 6.** ZINB model considering data stratified by Low Weeks (Model 3).

| Parameter |  | Model 3 (n=22,176) |  |  |  |
| --- | --- | --- | --- | --- | --- |
|  |  | IRR | 95% CI | z | p-value |
| Climate | Precipitation | 1.01 | [1.01-1.02] | 16.01 | <0.01 |
| Demographic | Income | 0.99 | [0.99-1.00] | -1.38 | 0.166 |
|  | Sex Ratio | 0.98 | [0.96-0.99] | -2.19 | 0.028 |
|  | Pop. Density | 1.00 | [1.00-1.00] | 5.67 | <0.01 |
|  | High Mob. | 1.26 | [1.12-1.42] | 3.85 | <0.01 |
| Environmental | Strat. Points | 1.03 | [1.02-1.03] | 12.69 | <0.01 |
|  | Apartment | 0.99 | [0.98-0.99] | -5.40 | <0.01 |
|  | Unpaved | 0.99 | [0.99-1.01] | -0.25 | 0.803 |
|  | Open Sewage | 1.02 | [1.00-1.03] | 3.47 | <0.01 |
| Scale | High Area | 8.14 | [3.57-18.56] | 4.99 | <0.01 |
| Year<br>Ref.: 2015 | 2007 | 0.16 | [0.13-0.19] | -23.27 | <0.01 |
|  | 2008 | 0.01 | [0.01-0.02] | -30.68 | <0.01 |
|  | 2009 | 0.00 | [0.00-0.01] | -26.45 | <0.01 |
|  | 2010 | 0.03 | [0.02-0.04] | -32.60 | <0.01 |
|  | 2011 | 0.04 | [0.03-0.05] | -31.72 | <0.01 |
|  | 2012 | 0.02 | [0.01-0.02] | -31.12 | <0.01 |
|  | 2013 | 0.09 | [0.08-0.18] | -27.63 | <0.01 |
|  | 2014 | 0.55 | [0.48-0.63] | -8.34 | <0.01 |

**Table 7.** ZINB model considering data stratified by High Areas (Model 4). * In 2009 there were no areas with high dengue incidence rates.

| Parameter |  | Model 4 (n=637) |  |  |  |
| --- | --- | --- | --- | --- | --- |
|  |  | IRR | 95% CI | z | p-value |
| Climate | Precipitation | 1.00 | [0.99-1.00] | 1.40 | 0.161 |
| Demographic | Income | 0.99 | [0.99-0.99] | -3.24 | <0.01 |
|  | Sex Ratio | 0.91 | [0.91-0.92] | -16.75 | <0.01 |
|  | Pop. Density | 1.00 | [1.00-1.00] | 5.62 | <0.01 |
|  | High Mob. | 1.69 | [1.52-1.88] | 9.91 | <0.01 |
| Environmental | Strat. Points | 1.02 | [1.02-1.02] | 9.37 | <0.01 |
|  | Apartment | 0.99 | [0.99-0.99] | -4.09 | <0.01 |
|  | Unpaved | 1.03 | [1.03-1.04] | 11.10 | <0.01 |
|  | Open Sewage | 0.98 | [0.97-0.99] | -5.27 | <0.01 |
| Scale | High Week | 2.22 | [1.72-2.88] | 6.10 | <0.01 |
| Year<br>Ref.: 2015 | 2007 | 0.29 | [0.21-0.41] | -7.22 | <0.01 |
|  | 2008 | 0.12 | [0.01-1.07] | -1.90 | 0.058 |
|  | 2009* | 1 | - | - | - |
|  | 2010 | 0.19 | [0.06-0.61] | -2.80 | <0.01 |
|  | 2011 | 0.26 | [0.16-0.45] | -4.98 | <0.01 |
|  | 2012 | 0.12 | [0.01-1.13] | -1.85 | 0.064 |
|  | 2013 | 0.24 | [0.17-0.35] | -7.64 | <0.01 |
|  | 2014 | 0.91 | [0.83-1.00] | -1.87 | 0.061 |

\* In 2009 there were no areas with high dengue incidence rates.

**Table 8.**
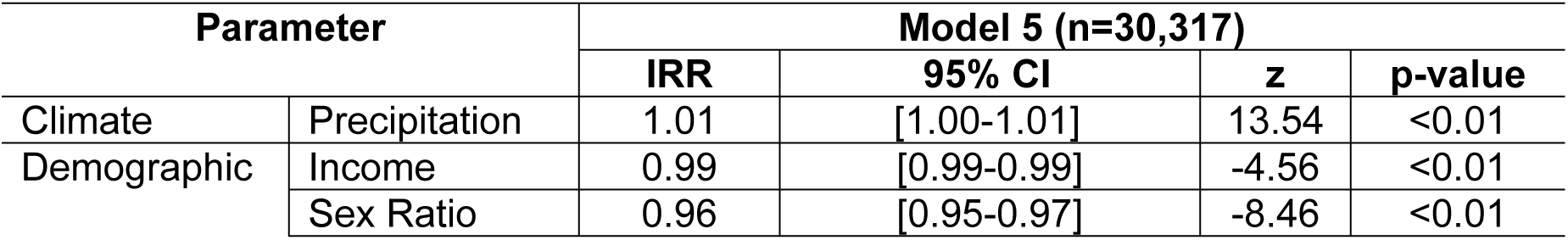
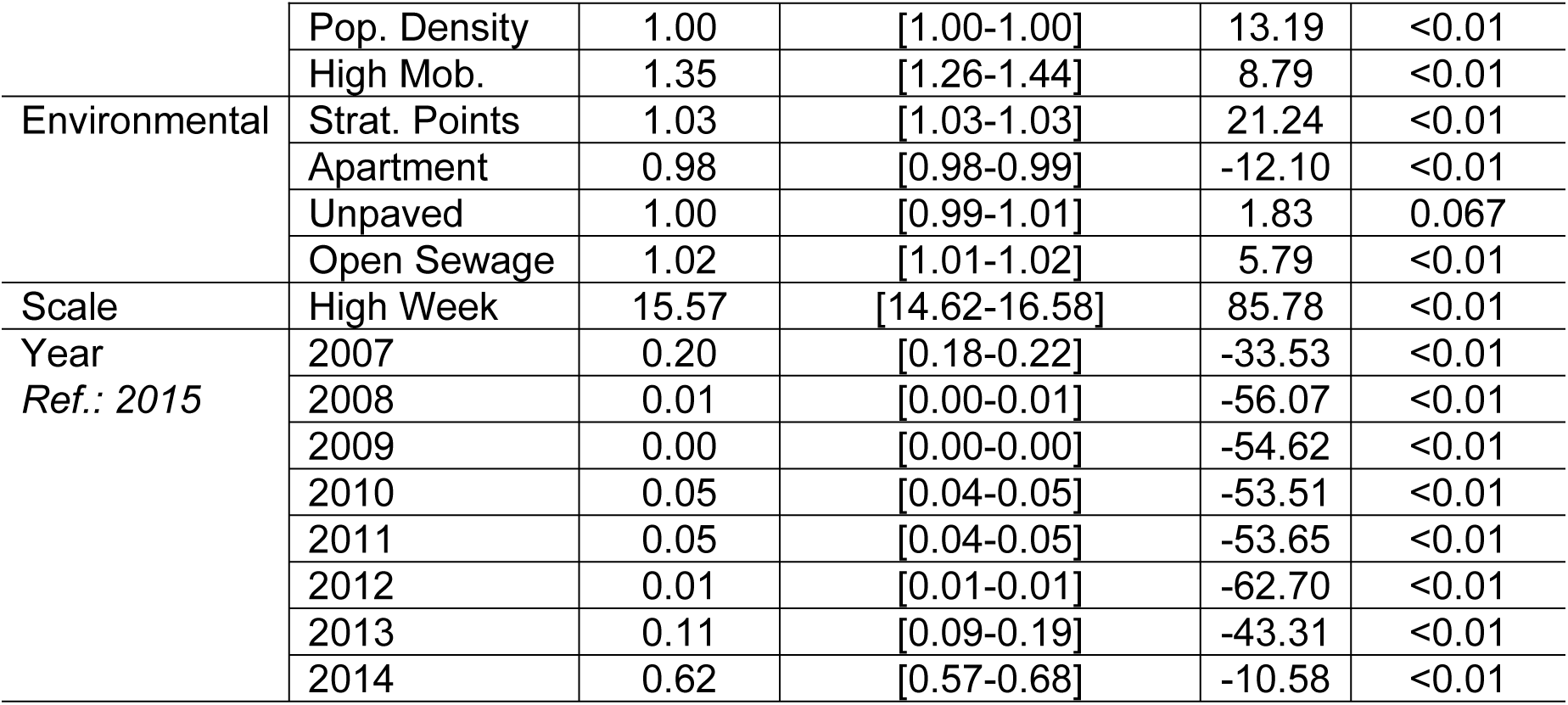
ZINB model considering data stratified by Low Areas (Model 5).

In Model 3, income per capita was protective against dengue fever, although not significant. Sex ratio was a protection factor against the disease in all stratified models, suggesting that more male residents are associated with a lower dengue incidence rate – in Models 2 and 4, the increase of one male per 100 female residents was accompanied by a decrease of 7% and 9% in the incidence rates, respectively. With respect to SPs, Models 1 to 5 showed that the addition of one SP contributed to an average increase of 3% in dengue incidence rates, with low variation across the different models. Similarly, the increase in 1% of population living in apartments reduced approximately 1% to 2% of dengue incidence rates on average across the distinct models’ stratification.

Population density did not differentiate residents across periods and areas with distinct levels of transmission (Models 1 to 5). Likewise, the increase in the proportion of households in each area with open sewage did not present substantial and consistent impacts on dengue incidence rates.

Living in a high incidence area increased dengue rates by 3 times in high transmission weeks (Model 2), while in low transmission weeks it increased rates by 8 times (Model 3). Conversely, during high transmission weeks living in high incidence areas increased dengue rates by 2 times (Model 4), while living in low incidence areas the dengue incidence was increased by more than 15 times (Model 5). Consequently, living in high transmission areas was associated with a higher increase in dengue incidence rates during low transmission weeks. On the other hand, high transmission weeks were most positively associated with increases in dengue incidence rates in low incidence areas.

## Discussion

This study aimed at investigating whether the daily pattern of population mobility was associated with DENV transmission in Campinas. Our results showed that population mobility was the most important determinant of dengue incidence, regardless of model specification.

While other studies have attempted to assess the role of population mobility, the geographical scale was often coarser [38–44]. Our study advanced current knowledge by showing that, at the local level, it was in the high incidence rate areas and during high transmission weeks that population mobility exerted the most expressive influence on DENV transmission. These results were consistent across different model specification, and after controlling for socioeconomic and environmental variables. While population mobility was a central element for DENV epidemics to occur, we found that high mobility and high dengue incidence rates did not perfectly overlap. For example, residents in the city center are highly mobile, but the area was not associated with high DENV transmission. Our results suggested that an explanation could be the type of housing, as the majority of the population in the city center live in apartments, often considered a protective factor against the DENV vector [45–47]. Residence in apartments was also a variable correlated with income in Campinas, as verticalization occurs more expressively in the city center, which is also the region with the highest concentration of more affluent people. This result corroborates previous findings of the role of income in DENV epidemics [26,48–53].

Although dengue seems to spread throughout the city, it is more prevalent in areas with precarious socio-environmental conditions, corroborating previous studies [13,54,55]. We found that DENV cases were more concentrated in – although not limited to – the South, Southwest and Northwest portions of the city. These are notable areas that concentrate less affluent populations and where environmental conditions favor the occurrence of *Aedes* breeding sites, highlighting the role of local inequalities in transmission. Unpaved streets and open sewage, proxies of local environmental quality, were significant risk factors for DENV in the high weeks and high areas transmission models (Models 2 and 4), corroborating previous findings [45,55–59]. With regards to population density, although some studies have found a positive association with dengue incidence [60,61], our results found no association, as high population density occurs also in the city center, where apartment buildings, a protective factor, predominates.

Precipitation (using an 8-week time lag) was a risk factor for DENV particularly during low weeks (May to January approximately) and low areas transmission models (Models 3 and 5, respectively). It is important to note that the most common types of breeding habitats in Campinas are recipients such as plant vases, animal drinkers, demountable swimming pools, cans, bottles and buckets, etc. [28]. Abundance of these containers are directly a result of human behavior and do not necessarily depend on rain water to be filled up. In addition, the effect of precipitation can happen in an indirect way. For example, the 2014 epidemics occurred during a severe drought in Campinas [21], which resulted in a behavior of storing water in buckets (not always properly covered) at home, favoring the proliferation of breeding habitats. There is still an open debate about the role played by sex ratio on dengue epidemics [63]. We found that, on average, men have fewer infections, particularly in high transmission weeks and areas (Models 2 and 4). Nevertheless, these results should be analyzed with caution. There is evidence that women usually seek health care assistance more frequently, not necessarily meaning that they are more affected by diseases, but, instead, that they tend to be more careful with their own wellness [64,65]. In Brazil, some studies found higher dengue notification among women [66,67], and others showed no significant sex difference [68,69]. In our data, from 2007 to 2015, 55% of patients that notified DENV cases were women while 45% were men, while the 2010 Population Census indicates that 52% of the residents in Campinas were women, and 48% were men [20]. This sex difference is important because there is a distinct mobility pattern between males and females. The OD Survey showed that women tended to have higher mobility within their area of residence, and therefore travel smaller distances than men (S2 Table). The extent to which this pattern implies lower exposure to an infection depends on the characteristics of the areas where they live and where they usually go.

Regarding Strategic Points, we found that the addition of one SP in an area tended to increase the dengue incidence rate by 3%, a result consistent with previous findings [26,27]. Although there are clear guidelines on how to monitor SPs, including mandatory inspection visits every fifteen days [24], limited financial resources, skilled professionals or even violence in an area can affect the regularity of these visits [26,54].

This study has some limitations. First, the used spatial unit of analysis was not the finest possible. However, given that the Origin-Destination Survey was conducted based on a sample of the population of each area, the spatial units could not be downsampled. Second, the flow of people traveling by air and by bus to other municipalities outside the Metropolitan Region was not addressed in this analysis. Yet, these travels represent a minor part in total daily travel, and thus are unlikely to change our results. Third, as it is the case for any administrative record on DENV infections, they capture only symptomatic cases (those that are registered by the health facility); that’s a limitation for any DENV study that relies on administrative records. Fourth, although reporting of DENV is mandatory, some private facilities do not fully report the occurrence of cases. Nevertheless, the private sector represents a minority of cases, unlikely to bias the results. Fifth, this study did not contemplate directions of population mobility in a more detailed time frame, restricting the concomitant analysis with the variation of DENV in space and time.

In this study, we analyzed a massive amount of DENV data, geocoded to patient’s residence addresses, in a detailed time-frame, for 9 consecutive years. Our results provided insights about DENV occurrence in a large urban center, suggesting patterns that could be similar in other large cities in Brazil, and also in Latin America. Most importantly, we shed light on the role that mobility plays on DENV transmission in a large city, which points to the need to improve surveillance mechanism to account for mobility patterns.

## Acknowledgments

We thank the Campinas Municipal Health Secretariat for providing part of the data used in this study.

## Supporting Information Legends

**S1 Dataset. Dengue incidence rates and covariates used in the study, by OD-weeks.** (Information will be available after paper approval).

**S1 Checklist: STROBE Checklist**

## Supporting Information

**S1 Table.**
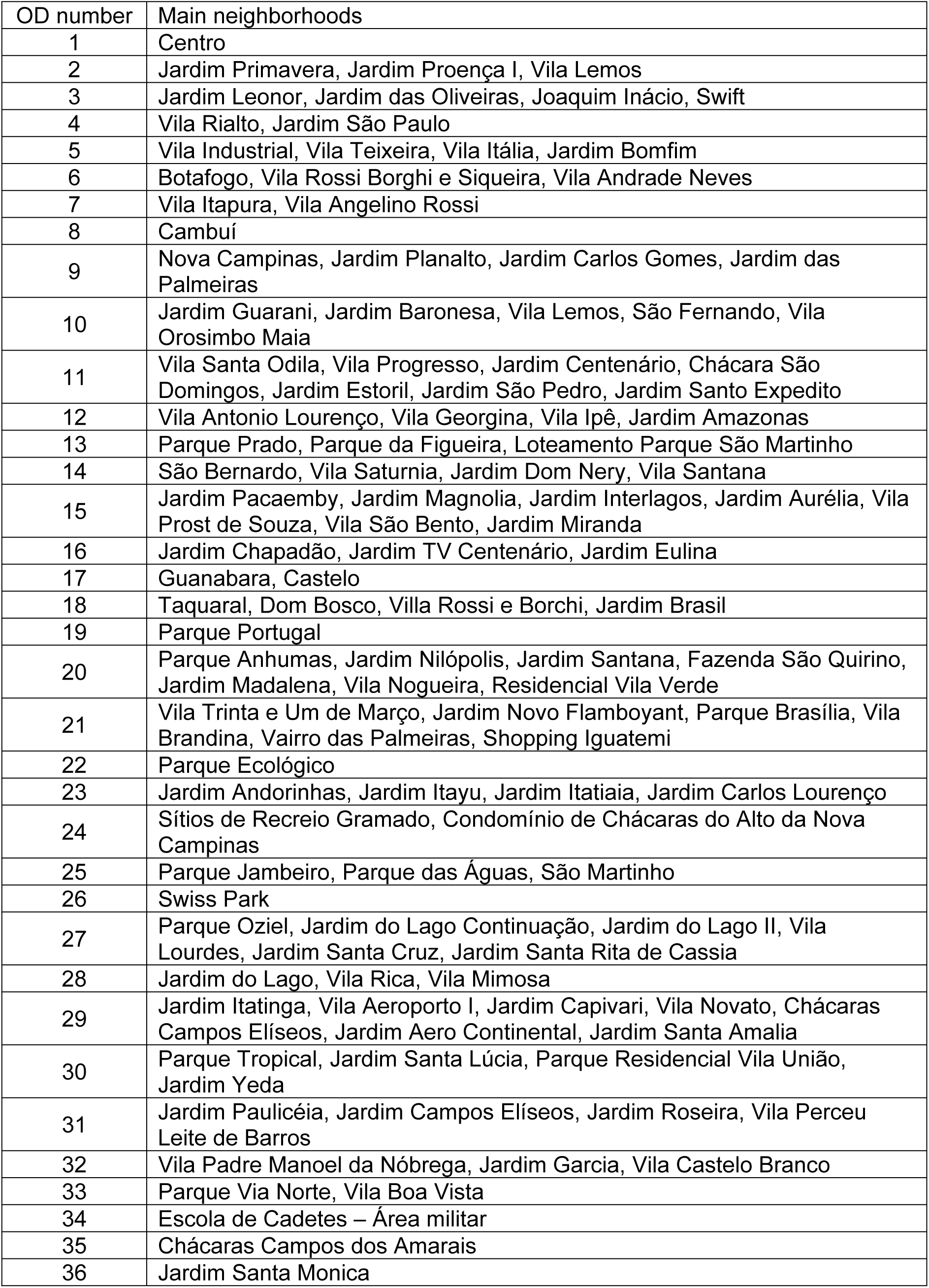
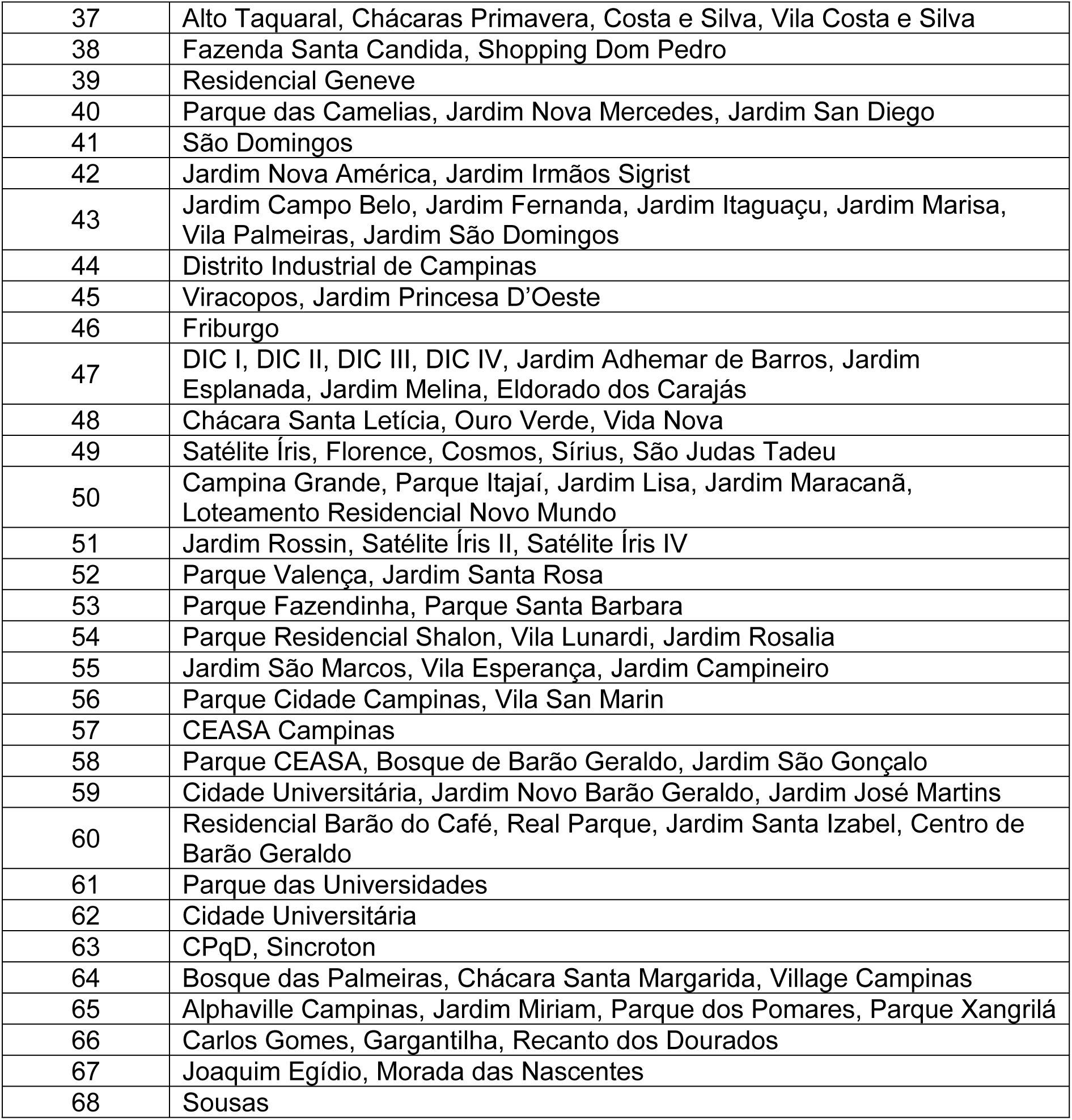
OD identification numbers and main neighborhoods comprised in each OD area in Campinas.

**S2 Table.** Total travels of Campinas inhabitants within the Metropolitan Region and the municipality by gender, Origin-Destination Survey, 2011.

|  |  | Female | Male | Total |
| --- | --- | --- | --- | --- |
| Total travels of Campinas inhabitants within the Metropolitan Region | Total Travels (%) | 1,031,636 (49.2) | 1,066,387 (50.8) | 2,098,023 (100) |
|  | Travels in resident area (%) | 294,750 (52.8) | 263,785 (47.2) | 558,535 (100) |
|  | Travels outside resident area (%) | 736,885 (47.9) | 802,602 (52.1) | 1,539,488 (100) |
| Total travels of Campinas inhabitants within the municipality | Total Travels (%) | 977,443 (50.6) | 952,438 (49.4) | 1,929,882 (100) |
|  | Travels in resident area (%) | 294,750 (52.8) | 263,785 (47.2) | 558,535 (100) |
|  | Travels outside resident area (%) | 682,693 (49.8) | 688,652 (50.2) | 1,371,346 (100) |

**S1 Fig.**
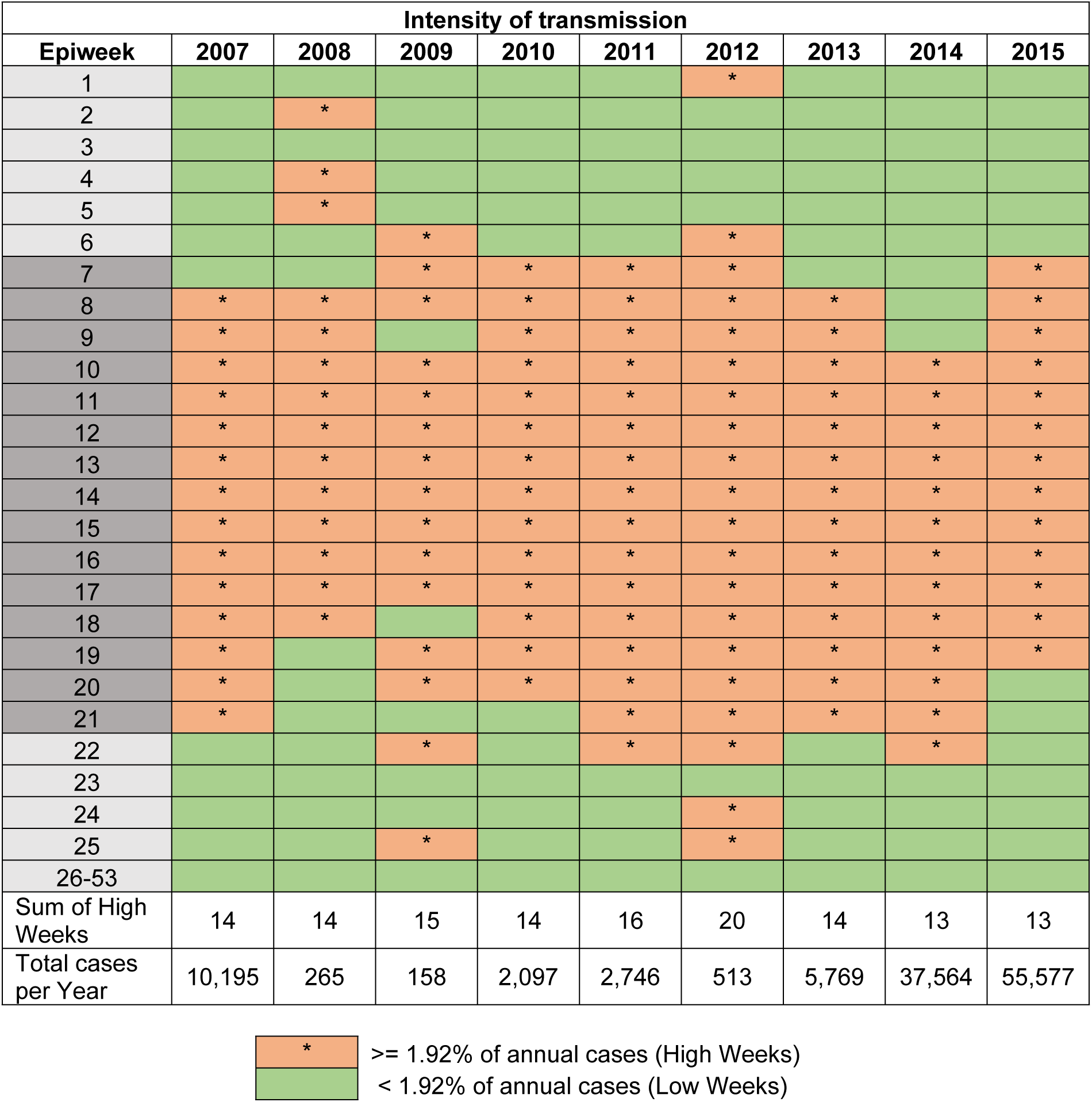
“High Weeks” - Classification of weeks according to the intensity of dengue transmission, Campinas, 2007-2015. Notes: 1-Only year 2014 had 53 epidemiological weeks. All the other years presented 52 EWs. 2-Weeks 7 to 21: dengue seasonality, with higher dengue cases concentration comprising the months February to May.

**S2 Fig.**
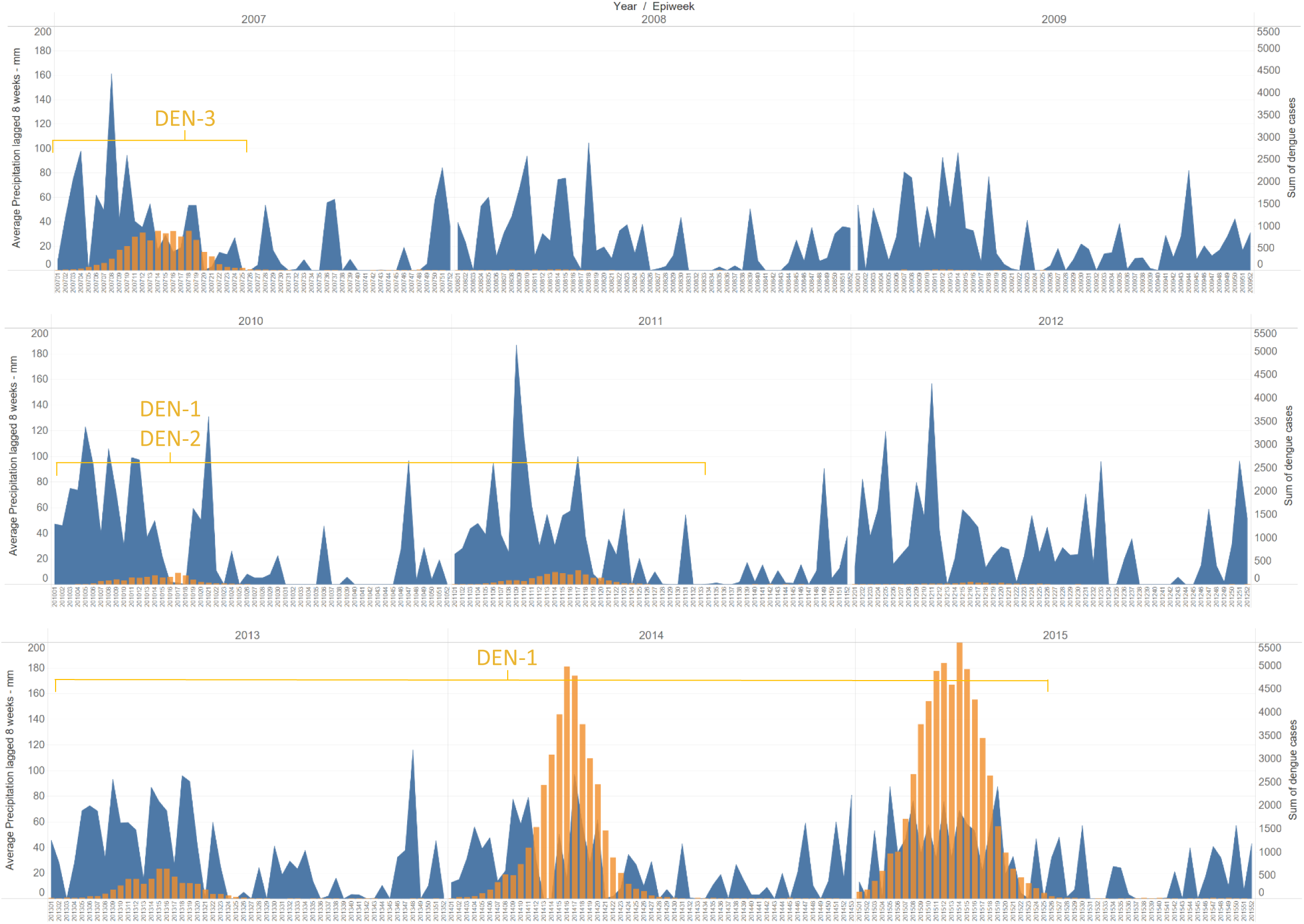
Sum of dengue cases and accumulated precipitation (lagged 8 weeks), by epidemiological weeks, Campinas, 2007-2015. Dengue cases represented in orange bars and precipitation in blue area. Predominant dengue serotype(s) indicated in yellow.

